# Waterbird communities in urban lakes: a comparison between Europe and South America

**DOI:** 10.64898/2025.12.24.696381

**Authors:** Lucas M. Leveau, Ianina N. Godoy, Maximiliano A. Cristaldi, Ornela V. Ganduglia

## Abstract

Freshwater environments can add environmental heterogeneity and biodiversity to cities. However, the relationship between the characteristics of urban lakes and their bird communities has been little studied, especially in the Global South. The objectives of this work are: 1) to compare bird communities between Europe and South America (SA); and 2) to analyze the relationship between bird communities and environmental characteristics such as lake size, surrounding vegetation and isolation from rural areas. We surveyed nine lakes in six SA cities of the Pampas Region and eight lakes in four European cities. The accumulated richness of birds was higher in SA. However, the accumulated richness of common species was similar between regions, while the richness of dominant species was higher in Europe. At the scale of each lake, total richness was positively related to the area of the lake. For the richness of common and dominant species, this relationship was maintained in Europe but was weak or null in SA. The results indicate that bird communities in urban lakes are a reflection of regional species assemblages. Lake size was essential for hosting a greater number of bird species, although its importance varied between regions.

## Introduction

Due to the expected rise in urban population, cities are likely to expand, impacting natural habitats [1]. Among these, freshwater habitats such as ponds and lakes are adversely affected by urbanization [2]. Urban growth harms freshwater habitats through destruction or habitat degradation, water pollution, flow changes, and invasion by non-native species [3,4]. On the other hand, ponds and lakes are created within cities for their aesthetic and recreational value [5,6]. However, their design and construction should also emphasize their role in promoting biodiversity conservation in urban areas [7], partly because of the positive value visitors can assign to biodiversity in urban freshwater habitats [8].

The impacts of urban ponds and lakes’ characteristics on biodiversity have mainly been studied in the Northern Hemisphere, focusing on amphibians, fish, and plants [7].

However, research on birds is much more limited. These studies have identified that several features, such as lake size, the cover of aquatic vegetation, water chemistry, and the extent of landscape urbanization, are important predictors of bird species diversity and composition [9–13]. For example, larger lakes and the presence of aquatic vegetation enhance bird diversity and support aquatic specialist bird species [10,13].

Additionally, the level of urbanization around lakes negatively affects bird diversity and the presence of aquatic specialist species [9,11]. Studies analyzing bird-habitat relationships in urban lakes of the Global South are much less common [14]. This disparity in knowledge between the global North and South limit our capacity to predict adequately the effects of urban lakés design on bird assemblages.

Therefore, this study aimed to compare bird communities in urban lakes across Europe and South America and to analyze differences in bird-habitat relationships between the continents. Due to dispersal processes from natural and rural areas into urban environments, we predict that species diversity and composition in each continent reflect their regional species pools. As a result, we expect higher species richness in South American urban lakes compared to those in Europe, given the greater aquatic species richness of the region [15–17]. Additionally, we anticipate that species diversity will be positively correlated with lake size and the percentage of aquatic vegetation cover in both continents. Lastly, we expect urban lakes near non-urban areas to support greater bird diversity due to increased dispersal opportunities and less surrounding urban cover.

## Methods

### Study area

The study was carried out in urban lakes located in parks of Europe and South America (Figure 1). According to regional bird guides and quantitative analyses [17–19], the two regions comprise uniform species pools of waterbird species. In Europe, bird surveys were conducted in eight urban lakes of four cities in three countries (Figure 1, Table S1). In South America, bird surveys were carried out in nine lakes of six cities located in the Pampean region of Argentina (Figure 1, Table S1). Lakes were separated by at least 200 meters. Most of lakes were surrounded by landscapes dominated (> 50% cover) by impervious surfaces (Table S2). European cities were colder (monthly annual mean = 12.70 °C) and drier (annual mean = 706.05 mm) than the South American cities (16.86 °C, 1009.08 mm) (Table S1). Most European cities had an Oceanic climate, whereas the South American cities had mostly a Subtropical humid climate (Table S1, [20]).

**Figure 1.**
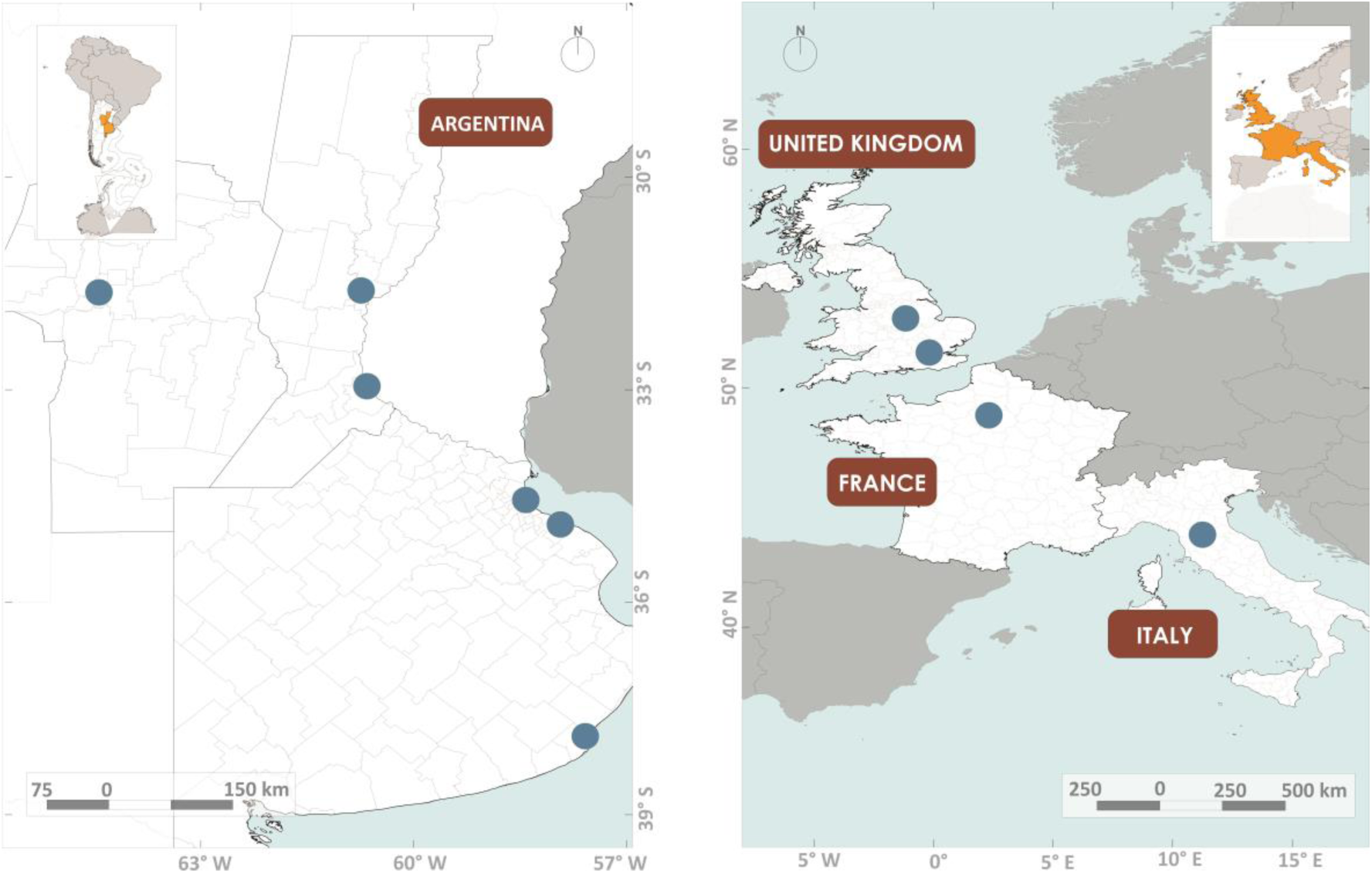
Location of cities surveyed in Europe (right) and South America (left). Map created with QGIS 3.8 Zanzibar, https://qgis.org/

### Bird surveys

Bird counts were carried out from points at the border of the lakes. The number of point counts varied between 1 and 7 according to the lake size (Table S2). Points were separated by at least 200 meters. Bird surveys were conducted under good weather conditions (no rain or strong winds) within the first four hours of the day mostly in spring of 2023 in South America and spring 2024 in Europe (Table S2). Birds were identified by sight or sound, and only those birds floating on water, drinking water, or perching on aquatic vegetation were included in the analysis. Obligate aquatic birds, such as wading birds, perched on trees at the edge of lakes, were also included. In South America, bird species introduced to urban lakes for aesthetic reasons, such as feral Greylag Goose (*Anser anser var. domesticus*) or feral Muscovy Duck (*Cairina moschata*), were not included in the analyses. In Europe, we included all bird species listed in the Collins Bird Guide [19].

### Environmental variables

Three environmental variables were measured in each lake (Table S2): area size (ha), percent cover of aquatic vegetation, and the minimum distance to rural areas (km).

Area size (mean = 4.71 ha, range = 0.05-21.64 ha) and the distance to rural areas (mean = 9.46 km, range = 0.95-18 km) were measured with Google Earth Pro. The aquatic vegetation cover (mean = 27.75 %, range = 0-80 %) was visually estimated at each point and included floating vegetation and coastal vegetation such as reeds or cattails.

## Data analysis

### Environmental variables

The difference of lake area between continents was tested with Student’s t-test using the t.test function in R [21]. Due to non-normal distribution of data (Kolmogorov test, P < 0.05), differences between continents of aquatic vegetation cover and the minimum distance to rural areas were tested with Mann-Whitney tests using the wilcox.test function in R [21].

### Bird diversity variables

We examined bird diversity using Hill numbers. Hill numbers represent the effective number of equally abundant species and vary based on their parameter q, which adjusts the weighting of species abundances [22]. Species richness (q = 0) assigns equal weight to all species, while Shannon diversity (q = 1) and inverse Simpson diversity (q = 2, hereafter Simpson diversity) assign increasing weight to species abundances. Therefore, Shannon diversity indicates the number of common species, whereas Simpson diversity reflects the number of dominant species [23]. The total number of species in each continent, according to each Hill number, was calculated using rarefaction curves generated by the iNEXT online software (chao.shinyapps.io/iNEXTOnline/) [24]. We used 84% confidence intervals for each curve, as described by MacGregor-Fors and Paiton [25], through 999 iterations. Curves with non-overlapping confidence intervals were deemed significantly different (P < 0.05). The sample coverage, which is the proportion of the total number of individuals that belong to the species detected in the sample [26], was estimated for each continent.

Bird diversity in each lake was calculated through Hill numbers, based on data of species abundances. The three Hill numbers were calculated using the species abundances for each lake with the hill_taxa function of the hillR package [27]. Hill numbers for each lake were analyzed in relation to environmental variables using generalized linear models (GLMs). For species richness (q = 0), the glm.nb function from the MASS package [28] was employed due to overdispersion. For Shannon and Simpson diversities, a Gaussian error structure was used. Models were obtained by backward elimination of non-significant variables (P > 0.05) from the full model using the anova function. Final models were compared with null models using a likelihood ratio test (LRT test) (p < 0.05). Multicollinearity among predictor variables was explored using the vif function of the regclass package [29]. The pseudo-rsquare of final models was obtained using the rsq function of the rsq package [30]. Model residuals distribution normality and heteroscedasticity were tested with the DHARMa package [31]. Final model results were plotted with the visreg package [32].

### Bird species composition

The relationship between bird species abundances and environmental variables was examined using distance-based Redundancy Analysis (dbRDA). This ordination technique employed species dissimilarity measures between lakes to link ordination axes with environmental variables [33]. We used Bray-Curtis dissimilarity, which considers species abundances, to compare lakes. dbRDA was performed with the capscale function from the vegan package [34]. Models were developed through backward variable selection and were compared to null models using a likelihood ratio test (LRT) (P < 0.05).

## Results

### Environmental variables

The area of lakes was similar between continents (t = 0.10, P = 0.921; Figure 2). The percent cover of aquatic vegetation tended to be higher in Europe than in South America (U = 16.5, P = 0.067; Figure 2). On the other hand, the minimum distance to rural areas was similar between continents (U = 68.5, P = 0.285; Figure 2).

**Figure 2.**
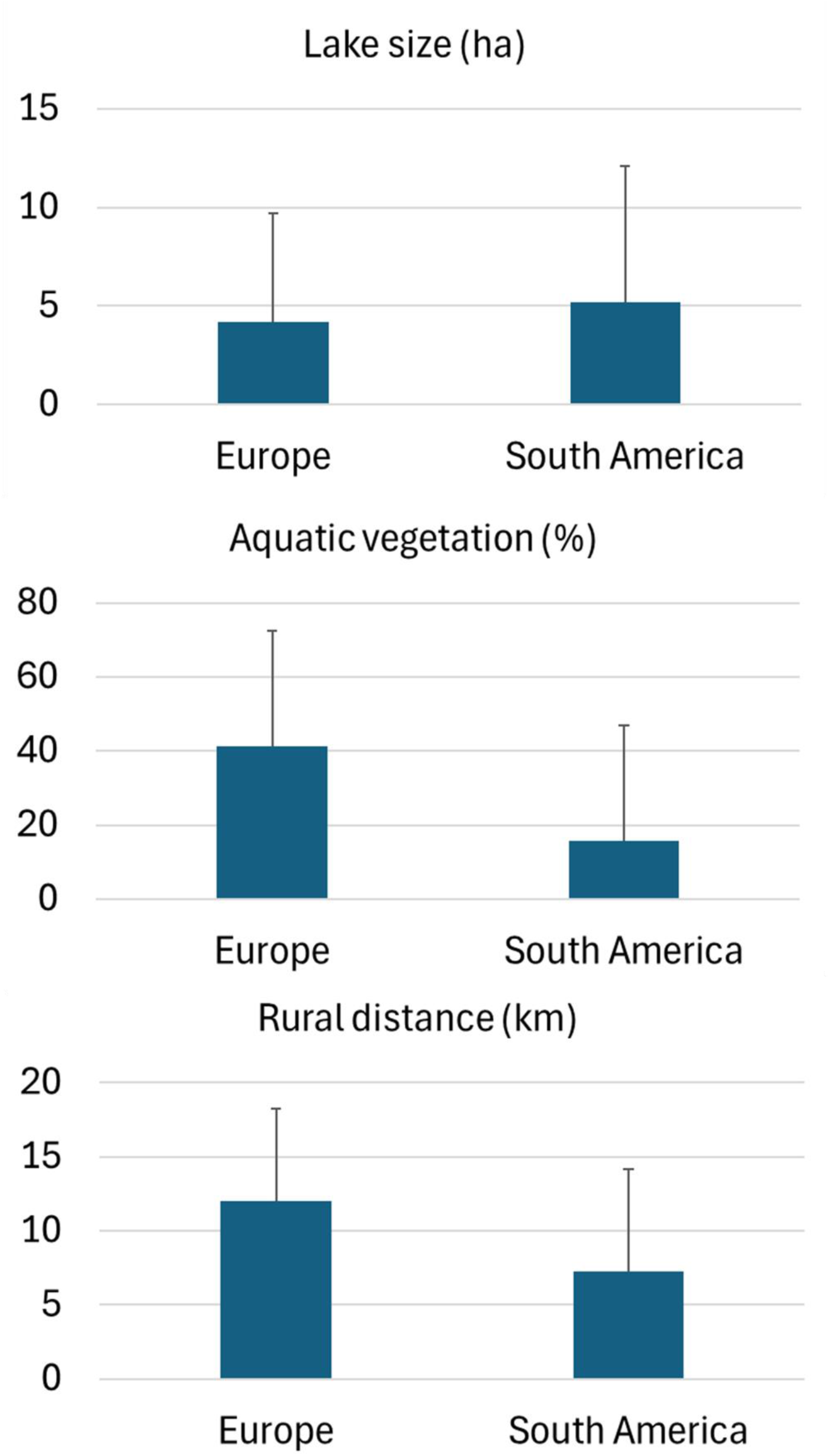
Habitat characteristics of urban lakes in Europe and South America. Bars indicate mean and vertical lines standard deviations.

### Bird assemblages

A total of 300 bird detections and 23 species were identified in Europe, while 805 bird detections and 40 species were identified in South America (Table 1). The most frequent and abundant species in Europe were the Eurasian Coot (*Fulica atra*), the Egyptian Goose (*Alopochen aegyptiaca*), and the Mallard (*Anas platyrhynchos*). In South America, the most frequent and abundant species were the Neotropic Cormorant (*Nannopterum brasilianum*), the White-winged Coot (*Fulica leucoptera*), and the Great Kiskadee (*Pitangus sulphuratus*). The number of introduced species was higher in Europe than in South American lakes (Table 1).

**Table 1.**
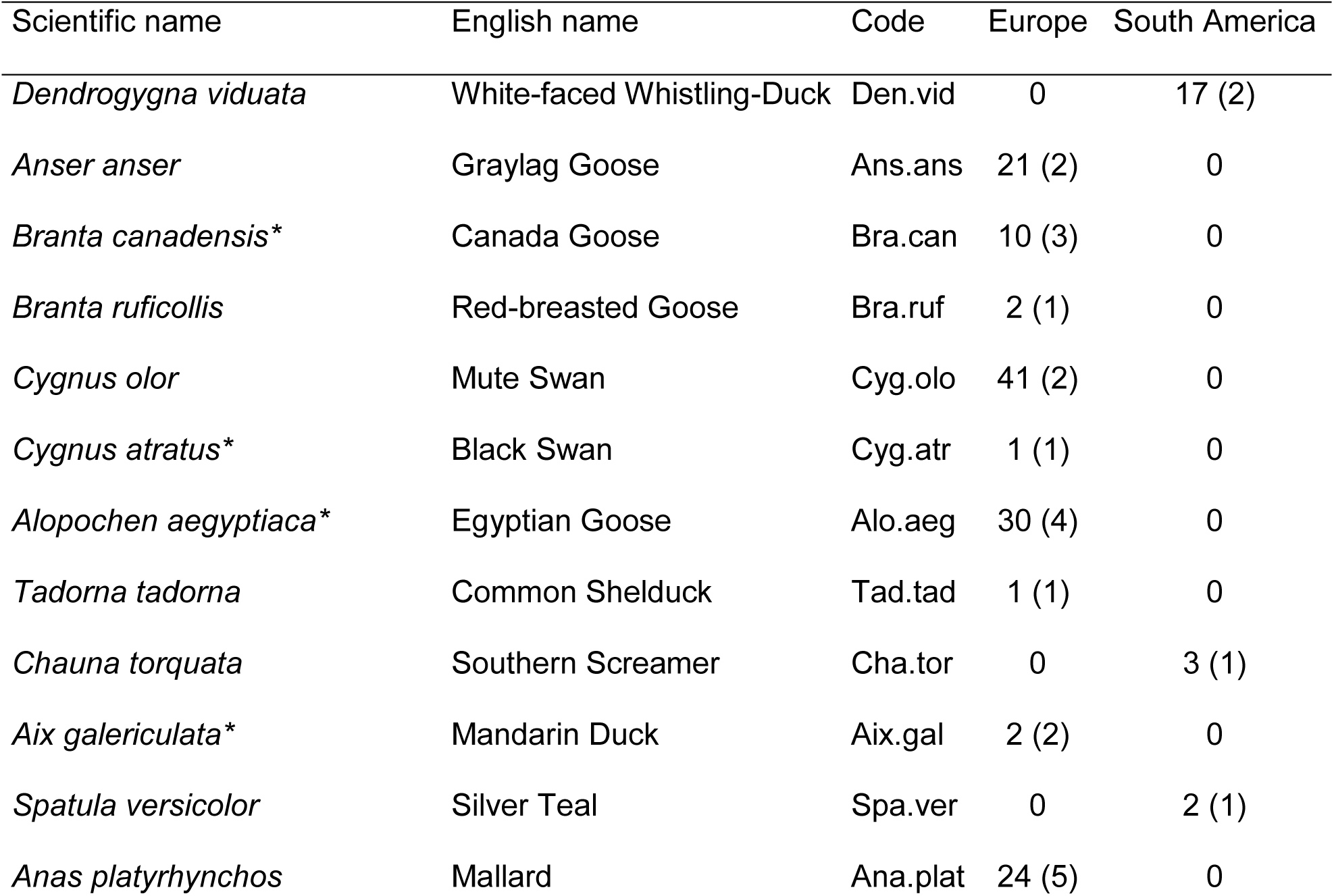
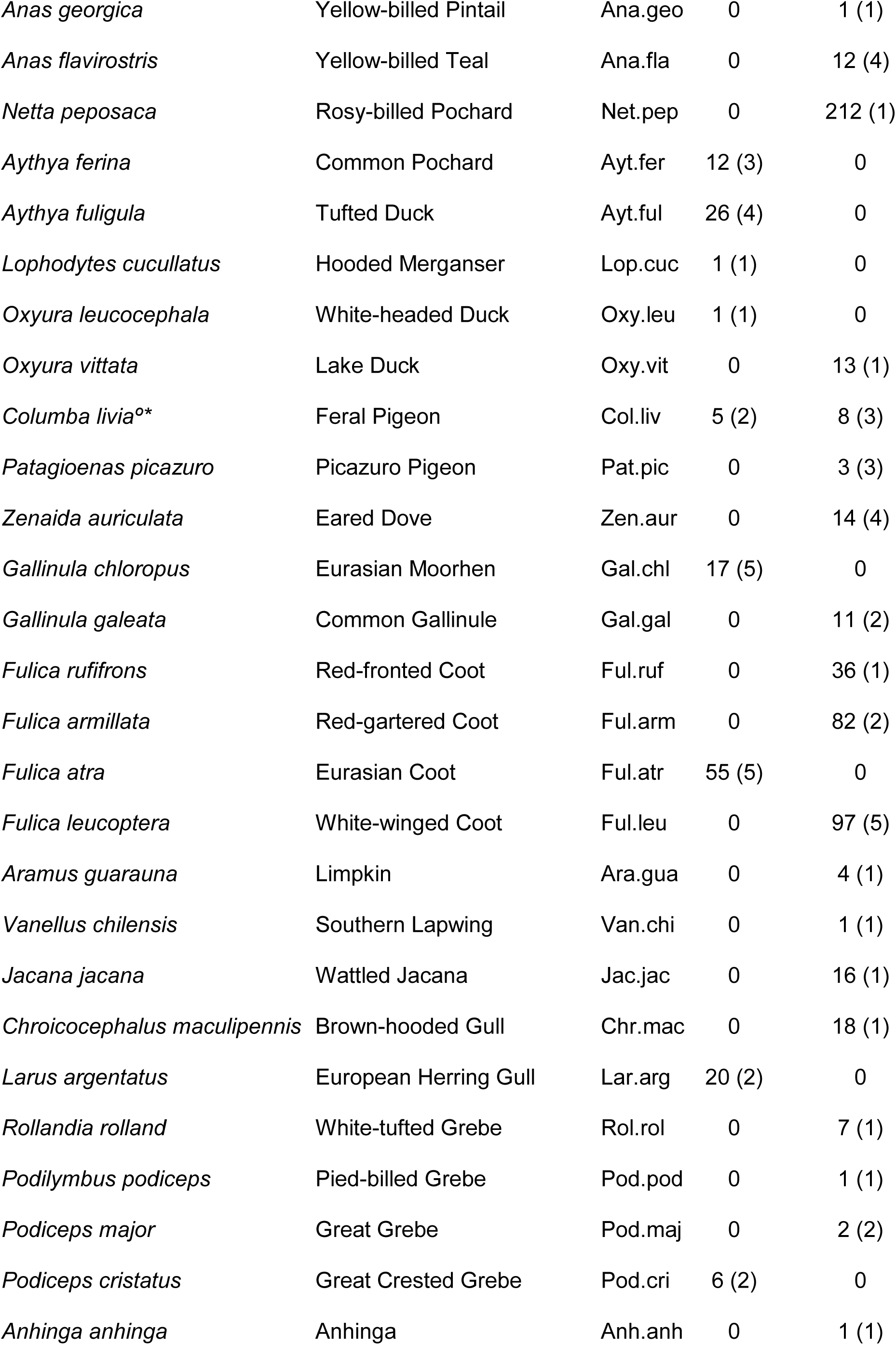
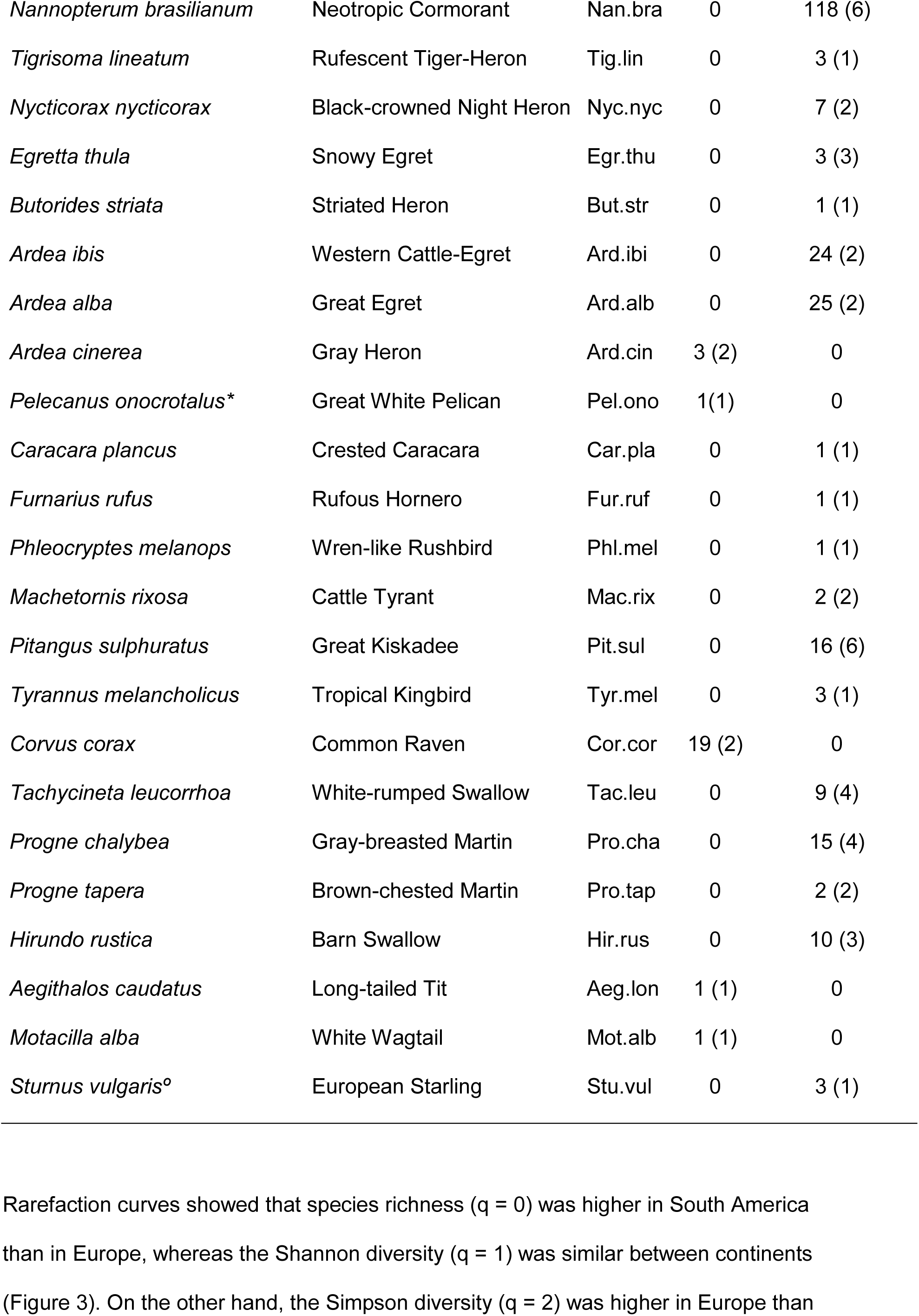
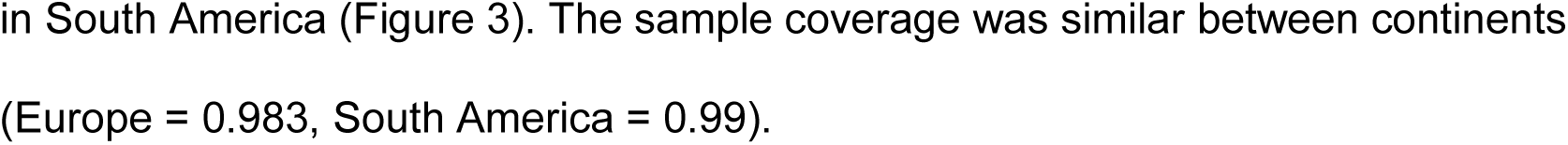
Bird species recorded in urban lakes of Europe (n = 8) and South America (n = 9) during spring. Numbers are species abundance and frequency (number of lakes, in brackets). Species codes issued in Figure 5 are given.*Indicates exotic species in Europe, whereas °indicates exotic species in South America.

**Figure 3.**
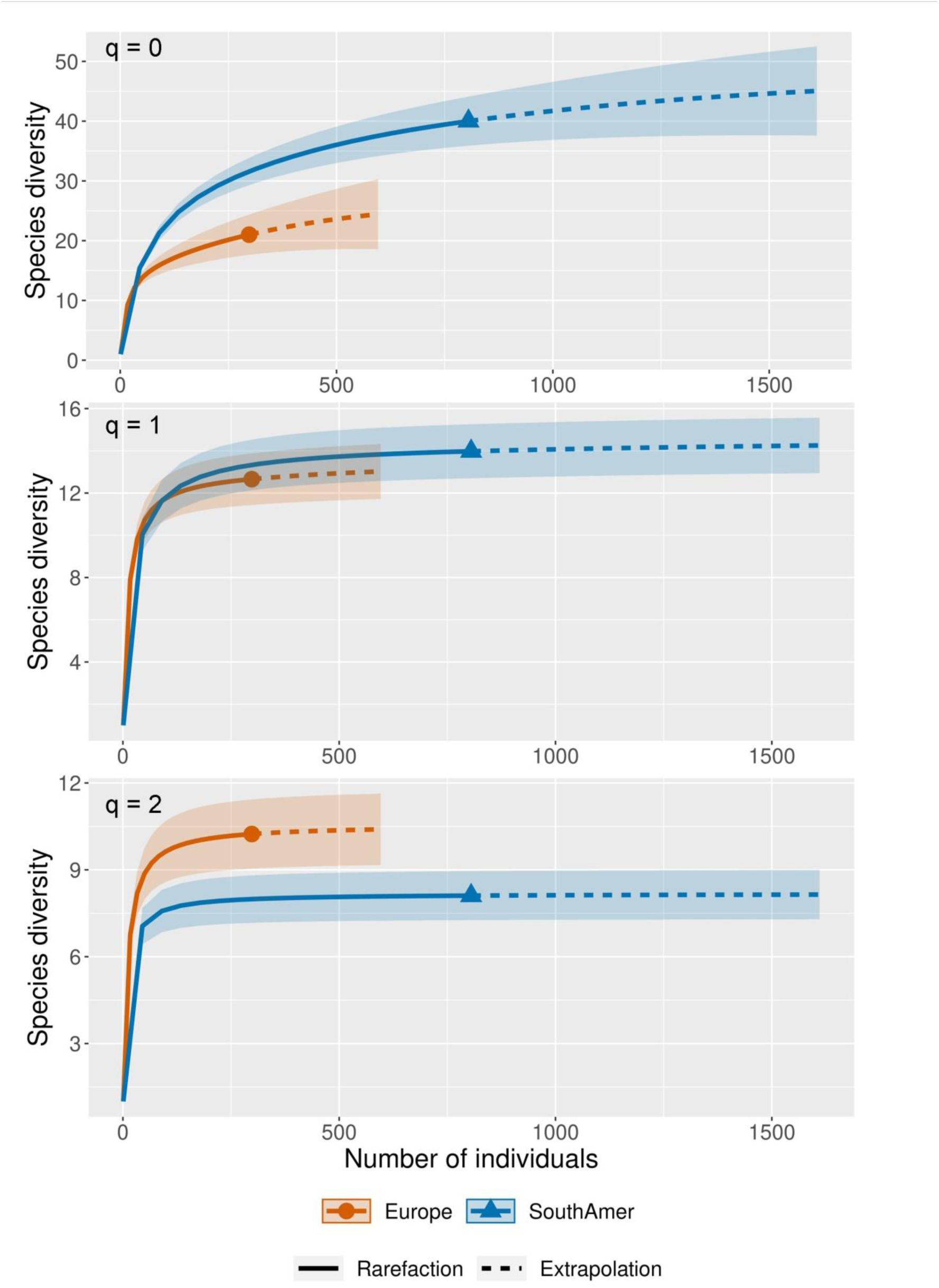
Rarefaction curves showing the species richness (q = 0), Shannon diversity (q = 1), and Simpson diversity (q = 2) in urban lakes of Europe and South America. Shaded bands show 84% confidence intervals.

The species richness per lake was positively related to lake size (LRT = 17.95; P < 0.001, r^2^ = 0.71; Table 2; Figure 4). The Shannon diversity and Simpson diversities related to lake size differently in Europe and South America (LRT = 65.27, P < 0.001, r^2^ = 0.55; LRT = 32.91, P < 0.001,r^2^ = 0.49; Table 2). Both diversities increased with lake size in Europe, whereas in South America the relationship was weaker or null (Figure 4).

**Figure 4.**
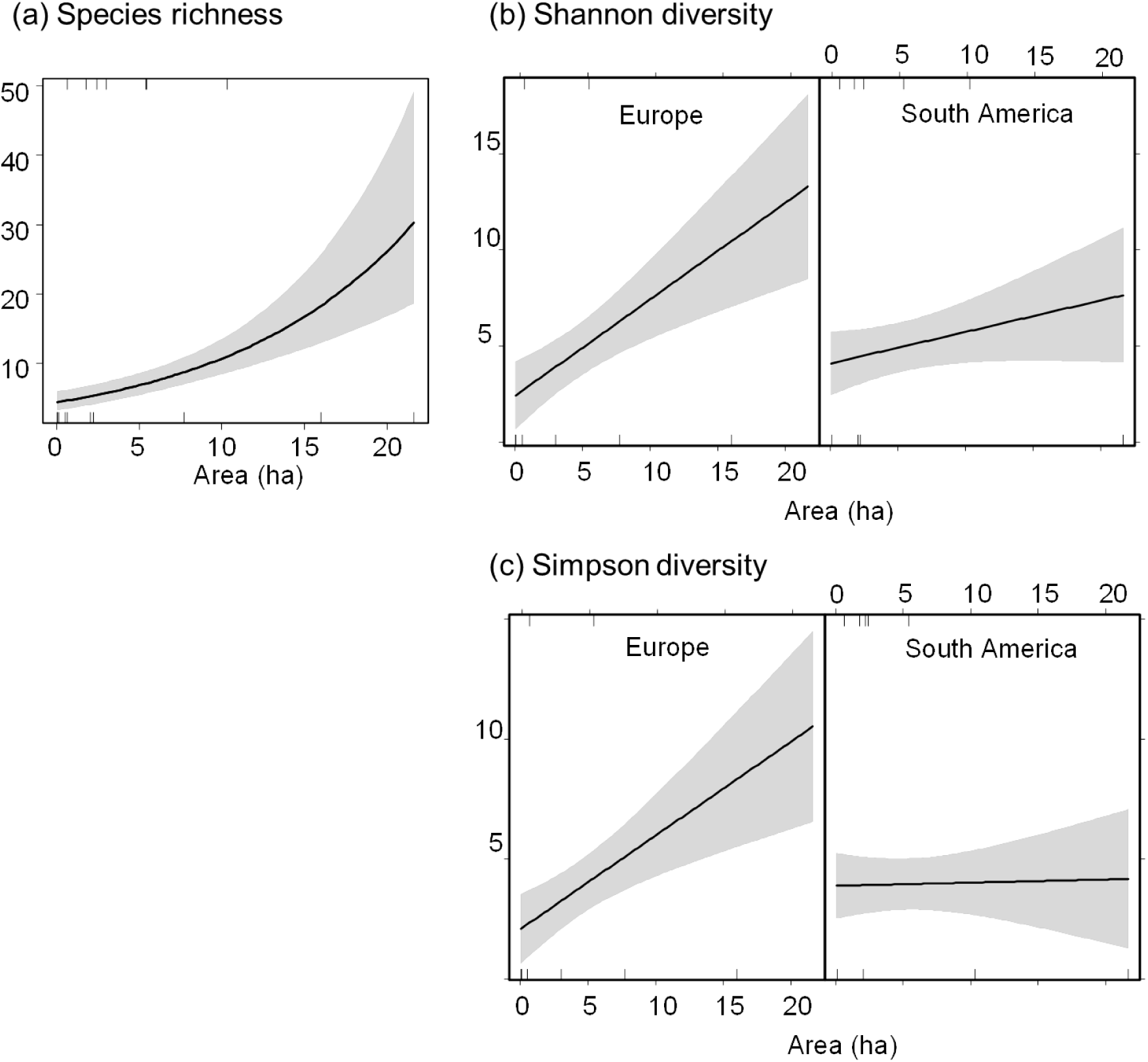
Relationship between a) species richness (q = 0), b) Shannon diversity (q = 1), c) Simpson diversity (q = 2) and environmental variables in urban lakes of Europe and South America. Black lines are fitted values and grey bands are 95% confidence intervals.

**Table 2.** Final generalized linear models showing the relationship between a) species richness (q = 0), b) Shannon diversity (q = 1), and c) Simpson diversity (q = 2) and environmental variables of urban lakes in Europe and South America.* Indicates interaction between variables. SE: standard error.

| Variable | Parameter | Estimate | SE | z value | P |
| --- | --- | --- | --- | --- | --- |
| a) Species richness ($q = 0$) | Intercept | 1.467 | 0.154 | 9.514 | <0.001 |
|  | Area (ha) | 0.090 | 0.015 | 5.921 | <0.001 |
| b) Shannon diversity ($q = 1$) | Intercept | 2.419 | 0.894 | 2.707 | 0.018 |
|  | South America | 1.680 | 1.226 | 1.371 | 0.194 |
|  | Area (ha) | 0.502 | 0.134 | 3.742 | 0.002 |
|  | South America*Area | -0.338 | 0.168 | -2.014 | 0.065 |
| c) Simpson diversity ($q = 2$) | Intercept | 2.088 | 0.739 | 2.826 | 0.014 |
|  | South America | 1.800 | 1.013 | 1.777 | 0.099 |
|  | Area (ha) | 0.389 | 0.111 | 3.510 | 0.004 |
|  | South America*Area | -0.376 | 0.139 | -2.711 | 0.018 |

Species composition was significantly related to continents and lake size (LRT = 3.21, P = 0.001, r^2^ = 0.31). Species characteristic of Europe were *Fulica atra* and *Cygnus olor*, whereas *Nannopterum brasilianum* and *Fulica leucoptera* characterized the South American lakes (Table 1; Figure 5). On the other hand, bigger lakes were positively related with the abundance of *Fulica leucoptera*, *Fulica armillata* and *Nannopterum brasilianum*.

**Figure 5.**
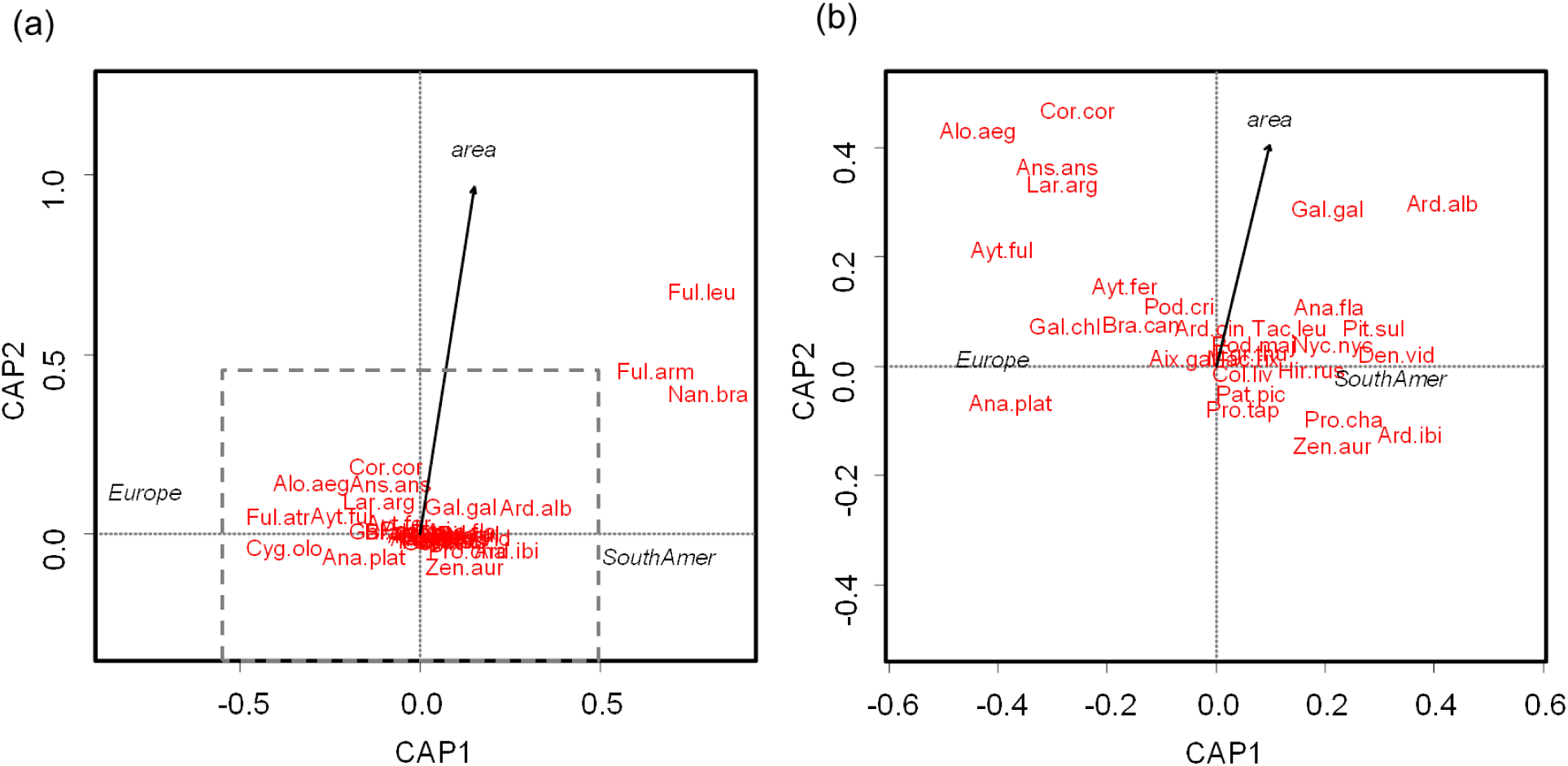
Distance-based redundancy analysis showing the relationship between environmental variables and bird species in urban lakes of Europe and South America. Figure b) shows the expanded area delimited by the dashed line in a).

## Discussion

The total bird species richness in urban lakes was higher in South America than in Europe, reflecting regional patterns of aquatic birds as shown by distributional maps [16,17]. However, this pattern reversed when we looked at the number of common and dominant species. On the other hand, the main predictor of bird species richness and species composition per lake was lake size, although its influence on the number of common and dominant species varied between continents.

### Patterns of bird species diversity

The total number of species was higher in South American urban lakes compared to those in Europe. This pattern aligns with other studies examining regional patterns of Anseriformes or Anatidae species based on distribution maps [16,17]. These studies have shown that the richness of aquatic species is mainly influenced by primary productivity and its seasonal changes [16,17]. Therefore, the higher temperatures, annual rainfall, and seasonal fluctuations in our South American study area likely increased the regional species pool, possibly helping their dispersal in urban lakes. Conversely, the total number of common species was similar across continents, and the number of dominant species was even higher in Europe than in South America. The increase in these bird diversity facets in Europe could be linked to local habitat features, such as the higher aquatic vegetation cover compared to South American lakes. Aquatic vegetation can improve the habitat diversity of urban lakes by offering nesting and food resources, as well as shelter for aquatic birds [14,35].

Bird species richness at the lake level was primarily influenced by lake size. This pattern aligns with studies in urban and agricultural landscapes [13,36,37]. Larger lakes can support more species due to increased and diverse resources [38]. However, the positive link between the number of common and dominant bird species and lake size was stronger in Europe than in South America. Europe experiences harsher climate and likely supports smaller bird populations compared to South America. The size of regional bird populations is a direct determinant of the rescue effect. This process impedes local population extinctions and the gradual loss of species with the decreasing size of the area [39]. Thus, due to smaller populations in Europe, the rescue effect is weaker than in South America, leading to a steady decline in species richness as lake area decreases. Conversely, the more favorable climate and larger regional bird populations in South America allow for a stronger rescue effect, making it more likely for urban lake populations to be saved from local extinction. As a result, the decrease or absence of decline in the numbers of common and dominant species with smaller lake sizes is observed in South America.

Aquatic vegetation cover and the minimum distance to rural areas did not show significant relationships with bird richness and diversity at the lake level. These results were unexpected, as several studies on birds and other taxa have indicated that these variables affect biodiversity in urban lakes [7,13]. Since these studies were conducted in a single city or region, and our research compared urban lakes across different continents, differences in spatial scale may explain the discrepancies between studies.

### Patterns of bird species composition

As expected, one of the top predictors of variation in bird species composition was the continent. The two regions studied, Europe and the Pampas in South America, consist of two different pools of bird species (for Anatidae species see Zeng et al. [17]), which later colonized the urban lakes. Europe had a greater diversity of exotic species that were not introduced to South America, thereby increasing the biotic differences between continents. Additionally, some aquatic cosmopolitan species that could be shared across continents, such as *Nycticorax nycticorax* [40], were only documented in South American urban lakes. Their presence in South American urban lakes likely reflects their higher population sizes in the region.

Lake area was also linked to bird species composition in lakes. The strongest association between bird species and lake size was observed in South America, where several coot species and the cormorant *Nannopterum brasilianum* were found in larger lakes. These types of lakes can support the coexistence of similar species such as *Fulica leucoptera* and *Fulica armillata*, which habitat use have been linked to different parts of lakes [41,42]. *Nannopterum brasilianum* has been associated with deep lakes that allow it to dive and fish for aquatic prey [43]. The lake area in our study was probably positively correlated with lake depth [44].

### Study limitations

Because of logistical constraints, the limited number of visits and surveyed lakes may restrict the strength of our results. Additionally, important environmental variables that can influence aquatic bird assemblages, such as water chemistry and human disturbance [4,7,45], were not measured in our study. We believe that collaboration between European and Latin American researchers can help overcome these limitations.

## Conclusions

The results indicated that bird species richness in urban lakes generally followed the macroecological patterns observed in natural areas. Urban lakes in South America exhibited higher species richness compared to those in Europe. However, when species abundances are taken into account, diversity values reveal an inverse pattern, being higher in Europe. Local-scale factors in European urban lakes, such as increased aquatic vegetation cover, may favor a higher number of common species compared to those in South America. Therefore, we recommend expanding the coverage of aquatic vegetation in the urban lakes of South America.

The size of the lake area was a fundamental predictor of species diversity and composition. However, our results indicated that the significance of lake size for the richness of common and dominant species was greater in Europe than in South America. The observed pattern may be linked to the scarcity of resources for birds in Europe, which could not sustain local populations of waterbirds in smaller lakes.

Therefore, designing larger urban lakes appears to be more necessary in Europe. Given the preliminary nature of this study, it is crucial to conduct more thorough analyses by increasing the sample size and the number of environmental variables. This goal could be accomplished by undertaking a collaborative study across continents or by utilizing citizen science data.

## Author contributions

Lucas M. Leveau: Conceptualization; Data curation; Formal analysis; Investigation; Methodology; Supervision; Software; Validation; Visualization; Writing – original draft; Writing – review & editing. Ianina N. Godoy: Investigation; Writing – review & editing. Maximiliano A. Cristaldi: Investigation; Writing – review & editing. Ornela V. Ganduglia: Formal analysis; Software; Writing – review & editing.

## Data availability

The datasets generated during and/or analyzed during the current study are available from the corresponding author on reasonable request.

