## Supplementary material for "Waterbird communities in urban lakes: a comparison between Europe and South America"

Table S1. List of cities surveyed in Europe and South America. Data provided are annual mean values of temperature, annual values of precipitation, climatic classification according to Köppen (1918), altitude above sea level, and human population according the last census.

| Continent | City | Temperature (ºC) | Precipitation (mm) | Climate type | Altitude (m.a.s.l.) | Population |
|---|---|---|---|---|---|---|
| Europe | Florence | 16.6 | 872.6 | Subtropical humid (Cfa) | 50 | 360930 |
| Europe | Paris | 12.8 | 634.3 | Oceanic (Cfb) | 33 | 10896433 |
| Europe | London | 11.3 | 601.7 | Oceanic (Cfb) | 15 | 14257962 |
| Europe | Nottingham | 10.1 | 715.6 | Oceanic (Cfb) | 61 | 328513 |
| South America | Buenos Aires | 18.25 | 1236.3 | Subtropical humid (Cfa) | 25 | 13395796 |
| South America | Santa Fe | 18.5 | 977 | Subtropical humid (Cfa) | 25 | 403878 |
| South America | Rosario | 17.8 | 1053.4 | Subtropical humid (Cfa) | 25 | 1348725 |
| South America | La Plata | 16.2 | 1072.7 | Subtropical humid (Cfa) | 26 | 787294 |
| South America | Mar del Plata | 14.0 | 946.1 | Oceanic (Cfb) | 38 | 682605 |
| South America | Alta Gracia | 16.4 | 769 | Subtropical humid (Cfa) | 557 | 60373 |

Table S2. Location and environmental characteristics of the urban lakes surveyed in Europe and South America. Data provided are date of bird surveys, geographical coordinates (latitude, longitude), lake area size (ha), percent cover of aquatic vegetation, minimum distance to rural areas (km), the number of point counts in each lake, and the urban level according to Leveau et al. (2022).

| Continent | City | Park | Lake | Date | Lat | Long | Area | Vegetation (%) | Rural dist | Points | Urban level |
|---|---|---|---|---|---|---|---|---|---|---|---|
| Europe | Florence | Fontana della Fortezza da Basso | Giardino della Forteza | 17/4/2024 | 43.782 | 11.252 | 0.51 | 15 | 0.95 | 1 | urban |
| Europe | Paris | Jardin des Tuleries | Exedre | 19/4/2024 | 48.863 | 2.327 | 0.05 | 70 | 18 | 1 | urban |
| Europe | Paris | Square du Temple - Elie Wiesel | Square du Temple - Elie Wiesel | 22/4/2024 | 48.864 | 2.361 | 0.06 | 80 | 17.65 | 1 | urban |
| Europe | London | The Regent´s Park | Boating lake | 25/4/2024 | 51.528 | -0.159 | 7.73 | 30 | 14 | 2 | urban |
| Europe | London | Hyde Park | Round Pound | 26/4/2024 | 51.506 | -0.183 | 3 | 0 | 13.2 | 1 | urban |
| Europe | London | Hyde Park | The Serpentine | 26/4/2024 | 51.505 | -0.168 | 16 | 10 | 14.4 | 2 | urban |
| Europe | London | St James´s Park | St James´s Park lake | 27/4/2024 | 51.502 | -0.135 | 5.44 | 70 | 14 | 3 | urban |
| Europe | Nottingham | Nottingham university campus | Jubilee pond | 10/4/2024 | 52.953 | -1.188 | 0.67 | 55 | 3.85 | 1 | suburban |
| South America | Buenos Aires | Reserva Costanera Sur | Reserva Costanera Sur | 21/10/2023 | -34.609 | -58.358 | 21.64 | 60 | 17 | 7 | urban |
| South America | Buenos Aires | Parque El Rosedal | El Rosedal | 13/11/2023 | -34.570 | -58.416 | 5.4 | 0 | 17 | 2 | urban |
| South America | Buenos Aires | Bosques de Palermo | Lago de Regatas | 7/11/2023 | -34.558 | -58.432 | 10.33 | 0 | 15 | 3 | urban |
| South America | Santa Fe | Parque Juan de Garay | Parque Juan de Garay | 28/10/2023 | -31.636 | -60.721 | 2.44 | 0 | 1.35 | 4 | urban |
| South America | Rosario | Parque Independencia | Parque Independencia | 1/10/2023 | -32.957 | -60.658 | 2.22 | 0 | 3.4 | 1 | urban |
| South America | Rosario | El Rosedal | El Rosedal | 1/10/2023 | -32.960 | -60.656 | 0.12 | 0 | 3.7 | 1 | urban |
| South America | La Plata | Parque Saavedra | Parque Saavedra | 25/10/2023 | -34.931 | -57.941 | 0.67 | 2 | 4 | 1 | urban |
| South America | Mar del Plata | Parque Camet | Laguna Camet | 16/10/2023 | -37.942 | -57.537 | 1.79 | 80 | 2.12 | 2 | periurban |
| South America | Alta Gracia | Tajamar | Lago Tajamar | 22/10/2022 | -31.656 | -64.435 | 2.05 | 0 | 1.46 | 1 | urban |
